# Development of nanobodies against human Survivin, applicable to suppress cell proliferation

**DOI:** 10.1101/2024.09.25.614874

**Authors:** Ayako Tagawa, Takeshi Fuchigami, Yusuke Miyanari

## Abstract

Survivin, a member of the inhibitor of apoptosis protein (IAP) family, is overexpressed in many cancers, making it a valuable target for therapy and a biomarker for cancer diagnostics. As Survivin contributes to cancer development by inhibiting apoptosis and promoting rapid cell division, inhibiting its activity is a promising strategy to overcome cancer resistance to apoptosis and reduce tumor growth. In this study, we developed nanobodies targeting human Survivin using M13 phage display and deep sequencing to identify high-affinity binders. Seven nanobody clones were selected and evaluated through ELISA, western blotting, and immunostaining. Clones 7 and 15, which specifically bind the alpha-helix region of Survivin, effectively suppressed cell proliferation by disrupting cell cycle regulation, likely by interfering interaction with intracellular binding partners. These findings suggest that Survivin-targeting nanobodies hold potential as therapeutic agents for cancer treatment, offering a novel approach to inhibiting tumor growth through the disruption of Survivin-mediated processes.

## Introduction

Survivin, a member of the inhibitor of apoptosis protein (IAP) family encoded by the BIRC5 gene, is expressed at low levels in most normal adult tissues, but is highly upregulated in various cancer types, making it both a potential target for cancer therapy and a valuable biomarker for cancer diagnostics (Jaiswal et al., 2015, Li et al., 2021). The elevated Survivin expression is associated with enhanced tumor survival, reduced efficacy of chemotherapy, and poor prognosis in patients, underscoring its significance in cancer biology (Lv et al., 2010). In normal adult tissues, Survivin expression is largely confined to proliferating cells, such as those found in the thymus, gastrointestinal epithelium, and during embryonic development (Sui et al., 2002). This restricted expression pattern makes Survivin a target for cancer diagnostics and therapeutics because targeting it selectively could minimize off-target effects on non-cancerous tissues. Survivin contributes to developing cancer by both inhibiting apoptosis and facilitating rapid cell division (Jaiswal et al., 2015). Furthermore, Survivin’s localization within the cell—both in the cytoplasm and nucleus—also plays a significant role in its function, with nuclear Survivin primarily involved in cell cycle regulation and cytoplasmic Survivin acting to inhibit apoptosis (Temme et al., 2003). Survivin is an essential component of the chromosomal passenger complex (CPC), a multi-protein assembly that governs key processes during mitosis, including chromosomal alignment, segregation, and cytokinesis (Vader et al., 2006). Within the CPC, Survivin acts as a scaffold that helps to recruit other proteins such as Aurora B kinase and INCENP (inner centromere protein) to the centromeres and midbody during cell division, ensuring proper chromosome segregation and the completion of cytokinesis (Honda et al., 2003). Survivin also functions as an inhibitor of apoptosis through interactions with other IAP family members and caspases, the key proteins responsible for executing apoptosis (Ambrosini et al., 1997). Specifically, Survivin directly inhibits caspase-9, a key initiator caspase in the intrinsic apoptotic pathway. By blocking caspase-9 activity, Survivin prevents the activation of downstream effector caspases such as caspase-3 and caspase-7, effectively halting the cell’s apoptotic machinery and promoting cell survival (Shin et al., 2001). This dual functionality makes Survivin a crucial target in cancer biology, where its overexpression allows tumor cells to evade apoptosis and continue proliferating.

Survivin’s multifunctionality is intimately tied to its unique structural features: the baculovirus IAP repeat (BIR) domain, the dimerization domain, and the alpha-helix domain (Fig. 1a, b) (Wheatley and Altieri, 2019). The BIR domain mediates protein-protein interactions with Caspase-9 and CPC proteins during mitosis, facilitating proper chromosomal segregation. The dimerization domain facilitates Survivin homodimer and also contributes to interaction with other CPC components, allowing it to localize to centromeres and the mitotic spindle during cell division. The alpha-helix domain is involved in interaction with proteins in the CPC, such as INCENP. The alpha-helix domain contributes to the precise localization of Survivin during mitosis, ensuring its correct function in chromosome alignment and cytokinesis.

**Fig. 1:**
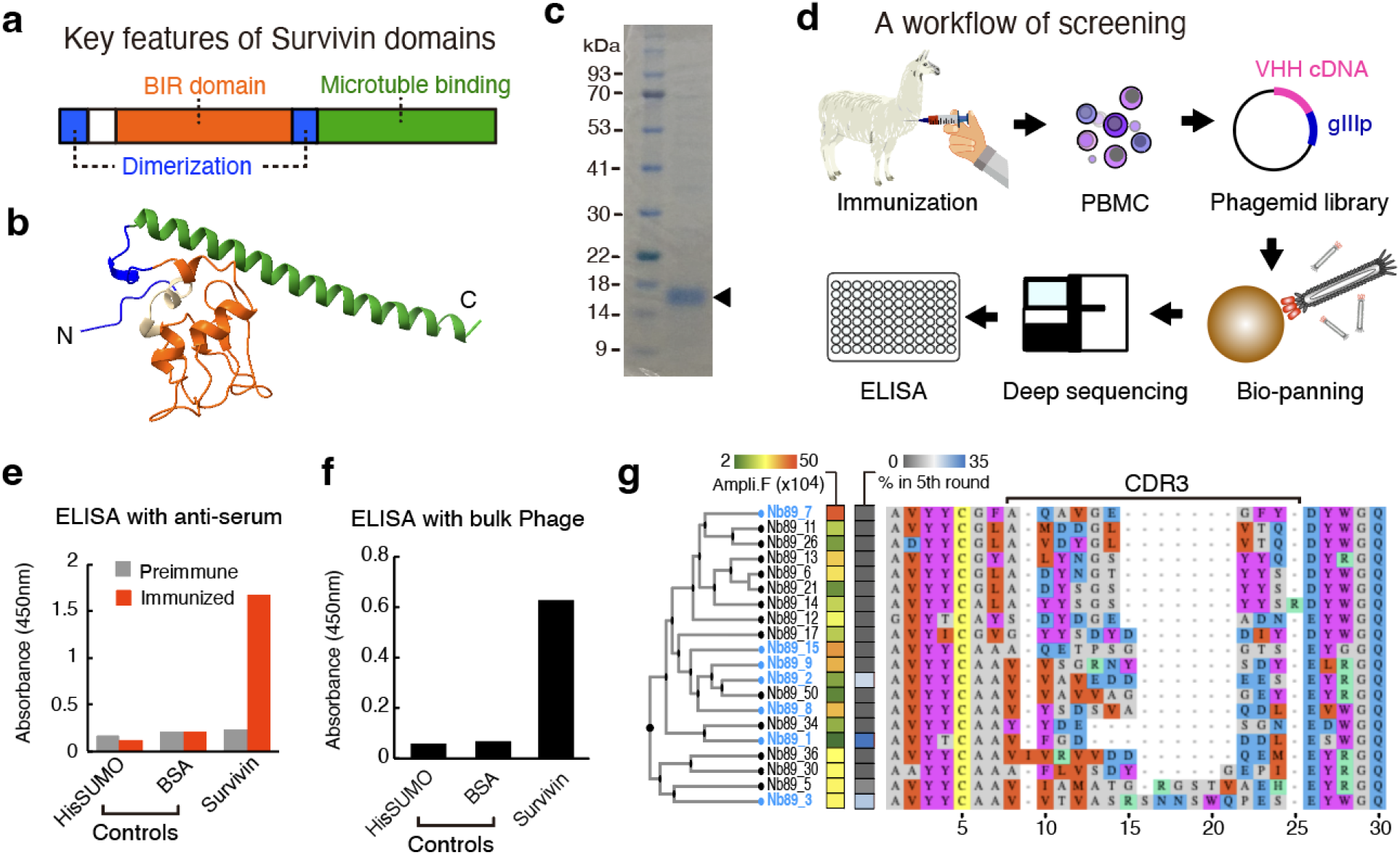
Screening of nanobodies against Survivin, a) Schematic illustration of Survivin with key functional domains. b) Structure of the Survivin protein. c) SDS-PAGE with CBB staining of Survivin protein used for immunization. d) Workflow of the screening process. e) ELISA using serums against the indicated proteins. f) ELISA with bulk phage from the final round of panning. g) Multiple alignment and clustering of amino acid sequences around the CDR3 region of the top 20 VHH candidates, ranked by amplification factor. The amplification factors and the proportion of each clone in the fifth round are shown. The clones selected for subsequent validations were highlighted in light blue.

The inhibition or downregulation of Survivin can disrupt critical mitotic processes, leading to defective cell division and eventually cell death. Efforts to inhibit Survivin expression or function using small molecules, antisense oligonucleotides, and immunotherapeutic approaches are actively being explored as promising strategies to overcome cancer’s resistance to apoptosis and reduce tumor growth (Li et al., 2019, Altieri, 2008, Olie et al., 2000). In this study, we developed nanobodies targeting human Survivin by employing a combination of bio-panning using the M13 phage display system and a deep sequencing strategy to identify specific high-affinity binders. The selected nanobody clones were evaluated across multiple applications, including ELISA, western blotting, and immunostaining, leading to the identification of two clones with superior functionality. Further investigation into the intracellular expression of these nanobodies revealed their ability to suppress Survivin activity, ultimately inhibiting cell proliferation in a tumor cell line. These findings suggest that Survivin-targeting nanobodies hold potential as a novel therapeutic approach for cancer treatment.

## Results

### Screening of nanobodies against Survivin

We have prepared a full-length Survivin protein expressed in *E*.*coli* and immunized it to a llama (Fig. 1c, d). After four rounds of immunizations, an ELISA assay using the snti-serum showed specific reactivity to the Survivin protein, in contrast to preimmune serum (Fig. 1e), indicating the establishment of a robust immune response against Survivin. To isolate specific binders to Survivin, we prepared a cDNA library of VHH genes from peripheral blood mononuclear cells (PBMCs) collected after the final immunization and performed bio-panning using the M13 phage display system with HisSUMO-Survivin protein as a bait. Since the bait was conjugated to magnetic beads or a 96-well plate by capturing HisSUMO, the native form of Survivin was expected to be preserved during the panning process. Following five rounds of panning, the enrichment of phages displaying anti-Survivin nanobodies was assessed by ELISA (Fig. 1f). The bulk phage specifically reacted to Survivin, confirming the successful enrichment of specific binders through panning. Given the elevated proportion of specific binders in the final library compared to the initial library, we performed deep sequencing of the VHH CDR3 regions, and calculated the amplification rate of each binder by comparing the library before and after panning. Following bioinformatic elimination of non-specific binders through subtraction with libraries against negative control baits, HisSUMO and magnetic beads, we obtained 35 clones that were significantly enriched in the final library. From these, we selected the top 20 clones, ranked by amplification factor (greater than 20,000), for clustering analysis of their CDR3 amino acid sequences (Fig. 1g). Highly diverse variations in CDR3 regions was observed, implying a broad range of binding specificities and affinity among the selected clones. We further selected 7 clones, consisting of the top 4 ranked by amplification factor and the top 3 with the highest proportion in the final library, for subsequent validations.

### Validation of nanobody clones

We expressed the selected 7 nanobodies as fusion proteins with an N-terminal HisNEDD8-StrepTagII in the SHuffle *E. coli* strain. The HisNEDD8-tag enhances protein solubility (Frey and Görlich, 2014a) while the SHuffle strain facilitates proper disulfide bond formation in the cytoplasm (Lobstein et al., 2012). The StrepTagII was used for detection. ELISA assays using lysates from *E. coli* expressing each clone demonstrated that all clones specifically reacted with the Survivin protein (Fig. 2a), validating our screening approach and confirming the functionality of these recombinant nanobodies. For further validations of these clones across several applications, we prepared tandem dimers of these nanobodies (Fig. 2b), with the expectation of enhancing their binding affinity as previously reported (Huang et al., 2021). The HisNEDD8-tag was cleaved from the nanobodies using NEDP1, a specific protease for NEDD8, to eliminate potential non-specific binding caused by the HisNEDD8-tag. For validation by Western blotting, we analyzed whole cell extracts from two cell lines: human HeLa cells and mouse embryonic stem cells (mESCs), both of which express Survivin (Cánovas, 2024, Guo et al., 2008). In addition, mESCs overexpressing FLAG-tagged human Survivin were prepared as a positive control. Western blotting showed that all clones detected both endogenous and overexpressed human Survivin, but not mouse Survivin (Fig. 2c), indicating their specificity to human Survivin, which shares 84.3% sequence identity with mouse Survivin. We next evaluated these clones by immunostaining mESCs transfected with plasmid DNA expressing FLAG-tagged human Survivin. As previously reported (Stauber et al., 2007), overexpressed Survivin was detected in both the cytoplasm and nucleus using an anti-FLAG antibody (Fig. 2d). Importantly, the fluorescent signals from all nanobody clones overlapped with those from the anti-FLAG antibody. Considering the western blot results, these findings suggest that all nanobodies can detect both denatured and native forms of the antigen. Notably, clones 7, 9, and 15 produced superior signals compared to the others. Next, we investigated whether these nanobodies could detect endogenous Survivin in HeLa cells by immunostaining. In contrast to its diffuse distribution across the cytoplasm and nucleus during overexpression, the subcellular localization of Survivin is dynamically regulated throughout the cell cycle, being restricted to specific regions such as the centrosome, mitotic spindle, midbody and centromeres (Temme et al., 2003, Uren et al., 2000).. Among the 7 clones, only clones 7, 9, and 15 produced specific signals in the expected subcellular regions, while the others showed no detectable signals (Fig. 2e). These results suggest that these three clones are suitable for detecting endogenous Survivin. Notably, centromeres were not stained by any of the three clones, in contrast to the anti-Survivin monoclonal IgG (Fig. 2f). This difference is likely due to variations in the antibody recognition regions of Survivin, as staining patterns of Survivin can vary depending on the antibody used (Fortugno et al., 2002, Wheatley et al., 2001).

**Fig. 2:**
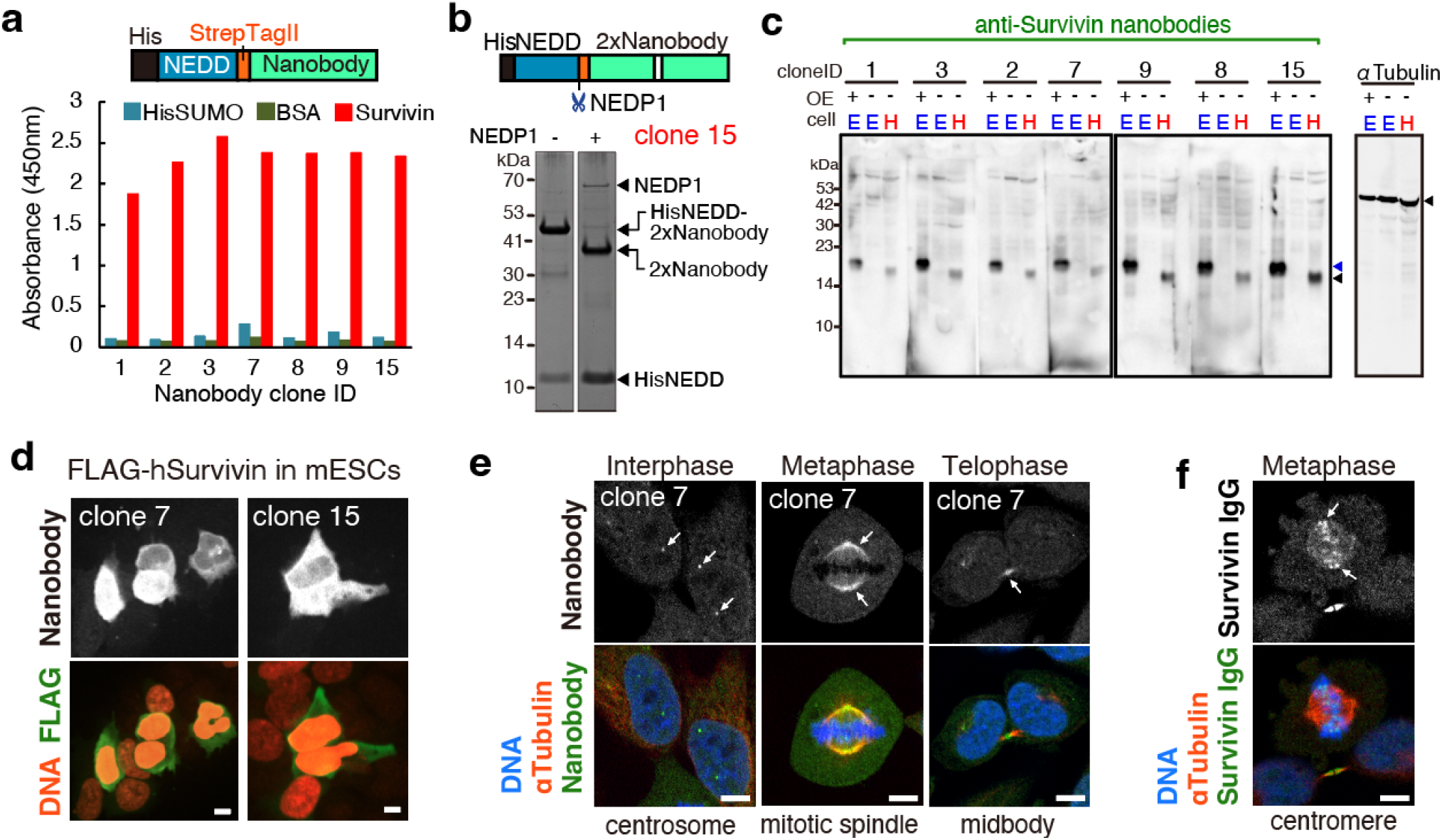
Validation of selected clones, **a)** Schematic illustration of the nanobody fused with HisNEDD8 and StrepTagII is shown at the top. ELISA results using *E. coli* lysates expressing selected nanobody clones against the indicated proteins are displayed. **b)** Schematic illustration of the tandem nanobody is shown at the top. The NEDD8 cleavage site by NEDP1 is indicated by a scissor icon. A representative image of SDS-PAGE with CBB staining for HisNEDD-tandem nanobody clone 15 with or without cleavage by NEDP1 is shown at the bottom. **c)** Western blot analysis using the indicated tandem nanobodies and anti-αTubulin IgG (as a loading control). Whole cell extracts from mESCs (E) and HeLa (H) cells were analyzed. OE: overexpression of FLAG-Survivin. Black and blue arrowheads indicate endogenous and FLAG-tagged Survivin, respectively. **d)** Representative images of fluorescent immunostaining of mESCs transfected with pFLAG-Survivin. Nanobody (gray), FLAG (green), and DNA (red, stained with PI) are shown. Representative images from immunostaining tests using the 7 nanobody clones are shown for clones 7 and 15. **e)** Representative images for immunostaining of endogenous Survivin in HeLa cells detected using the nanobody clone 7, along with α-Tubulin antibody and DNA (DAPI) staining. Cell cycles of cells in each image are indicated on the top. The subcellular localizations indicated by the arrows in each panel are shown below the corresponding panel. Images using clone 7 are shown as representatives. **f**) Representative images for immunostaining of endogenous Survivin in HeLa using anti-Survivin monoclonal antibody.

### Characterization of selected nanobodies

Given their superior performance in immunostaining, we selected clones 7, 9, and 15 for further characterization of their antibody recognition regions. We first prepared three truncated Survivin proteins (Ag1 to Ag3), each containing characteristic domains of Survivin (Fig. 3a), and analyzed them by Western blotting using three nanobody clones (Fig. 3b). We found that all these clones reacted to Ag3 but not Ag1 or Ag2, indicating they recognize the alpha-helix region. In contrast, the anti-Survivin IgG antibody used for immunofluorescence in Fig. 2f recognized Ag1 and Ag2, suggesting it binds to the BIR domain. To determine the more precise regions recognized by these nanobodies, we predicted the structures of the Survivin-nanobody complexes using AlphaFold-Multimer(Jumper et al., 2021; Evans et al., 2021, Bryant et al., 2022). Consistent with the Western blotting results using truncated proteins, all clones were predicted to bind the α-helix region (Fig. 3c, 3d), though their binding orientations and predicted epitopes differed from one another. Importantly, all predicted epitopes included amino acid sequences that differ from those in mouse Survivin, explaining the nanobodies’ specificity to human Survivin. Surface plasmon resonance assays for the monomeric nanobodies of clones 7 and 15 revealed binding affinities, with dissociation constant (Kd) of 1.52 µmol/L and 16.1 µmol/L, respectively.

**Fig. 3:**
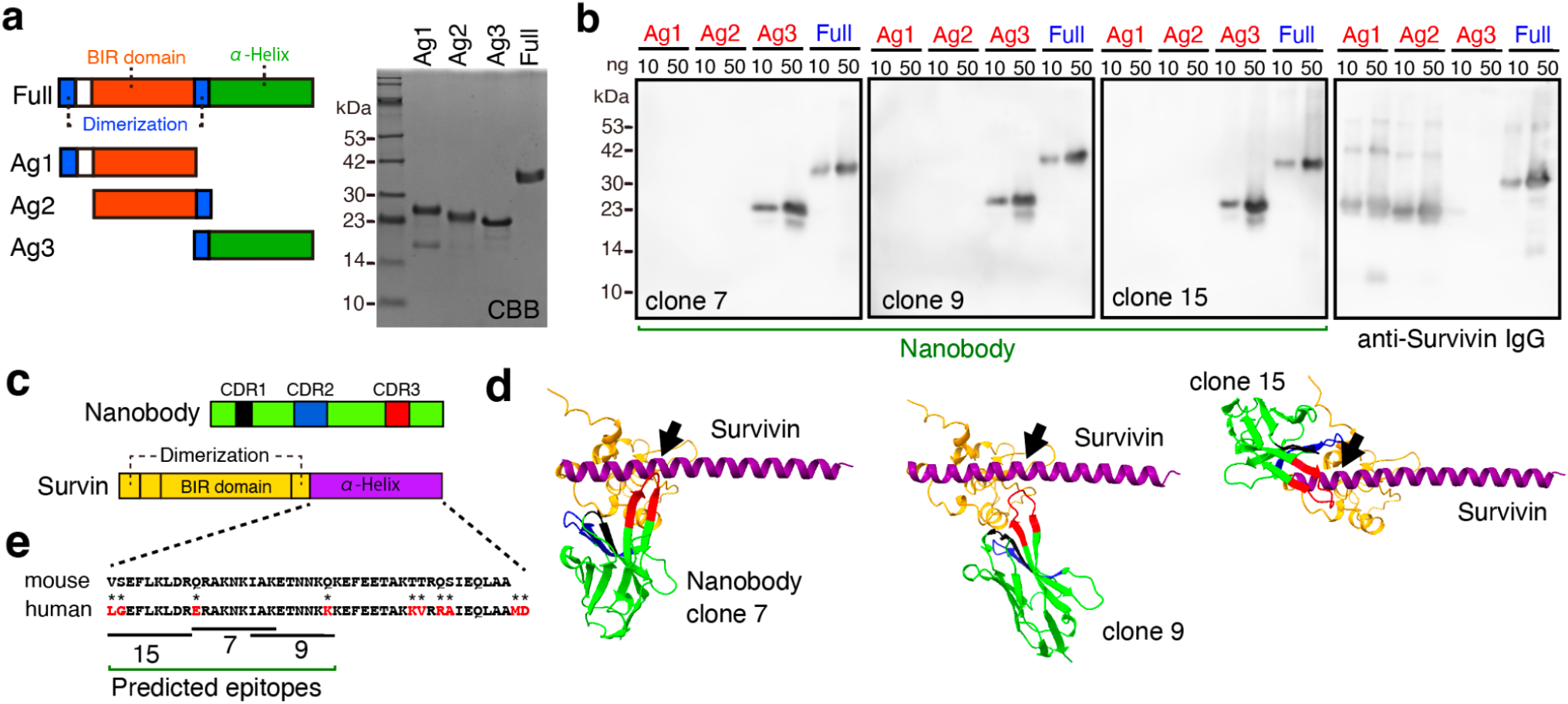
Characterization of nanobodies. **a)** Schematic illustrations of truncated Survivin proteins and their SDS-PAGE analysis with CBB staining. **b)** Western blot images showing the analysis of 10 and 50 ng/well of the indicated proteins using nanobody clones 7, 9, and 15. **c)** Key features of Survivin and nanobody. The color of each domain corresponds to the colors shown in (b). **d)** AlphaFold2 predicted structures of the Survivin-nanobody complexes. The binding sites were indicated with arrows. **e)** Amino acid sequences of α-helix of human and mouse Survivin, with asterisks on different amino acids. The predicted epitopes of indicated clones are marked with underlines.

### Suppression of Survivin activity by intracellular expression of nanobodies

Since Survivin is highly expressed in cancer cells and plays dual roles in promoting cell proliferation and preventing apoptosis (Garg et al., 2016; Wheatley and Mcneish, 2005), one potential application of nanobodies is the inhibition of Survivin. To evaluate the effects of intracellular nanobody expression, we used a lentiviral system to express anti-Survivin nanobodies from clones 7 and 15, as well as an anti-MoonTag nanobody as a control. We first assessed the impact on HeLa cell proliferation by counting viable cells 4 days post-infection and found that the expression of clone 7, but not clone 15, led to a substantial and statistically significant reduction in cell number (Fig. 4a). This result suggests that clone 7 possesses intracellular activity that suppresses Survivin’s function. Western blotting for the FLAG-tagged nanobodies confirmed their expression, although clone 15 showed relatively lower expression levels compared to the others (Fig. 4a, bottom). To further investigate the effects of the nanobodies, we analyzed their impact on the cell cycle by quantifying DNA content using flow cytometry (Fig. 4b). The results showed that clone 7, but not clone 15, caused a significant decrease in the G2/M phase population, accompanied by an increase in the G1 phase. We also assessed the effect on apoptosis using Annexin-V staining. Expression of clone 7 resulted in a slight increase in the proportion of apoptotic cells compared to the control, although this difference was not statistically significant (Fig. 4c). TThese findings suggest that clone 7 primarily suppresses cell proliferation by affecting the cell cycle. The superior effect of clone 7 over clone 15 may be attributed to its higher expression level and stronger binding affinity. To further explore their effects on cell proliferation, we overexpressed nanobodies using plasmid DNA, which leads to higher expression levels than the lentiviral system. In this approach, both clones exhibited a reduction in cell number, surpassing the effects observed with the lentiviral system (Fig. 4d). Therefore, we concluded that both clones possess inhibitory effects on Survivin when expressed at sufficient levels.

**Fig. 4:**
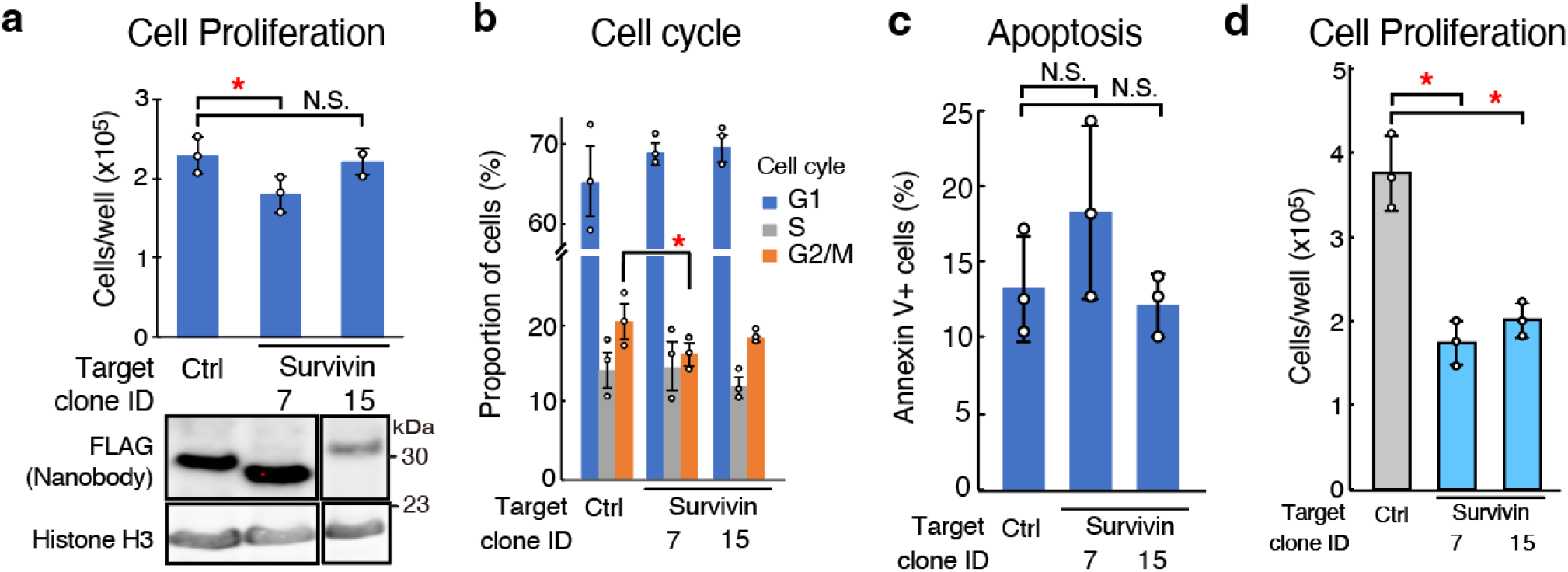
Effects of intracellular expression of anti-Survivin nanobodies. **a)** HeLa cells were infected with lentiviruses encoding the indicated nanobodies fused with an N-terminal FLAG tag. Viable cell counts 4 days post-infection are shown at the top, with Western blot analysis of the indicated proteins at the bottom. **b**) Cell cycle analysis was determined by flow cytometry using DAPI staining (n=3). **c**) Proportion of cells positive for Annexin V staining (n=3). **d**) The indicated nanobodies were ectopically overexpressed by plasmid DNA transfection. Student’s t-tests (one-sided, unpaired, n=3) were performed. * denotes p < 0.05, and N.S. indicates not significant.

## Discussion

In this study, we successfully identified nanobodies targeting human Survivin through a combination of M13 phage display biopanning and deep sequencing strategies. Our findings demonstrate the effectiveness of deep sequencing in nanobody screening and selection, as well as the functional potential of the identified clones in various applications. Our results provide insights into the suitability of specific nanobodies for downstream applications including ELISA, western blotting, and immunostaining. Notably, we identified two nanobody clones which can suppress cell proliferation assays through intracellular expression, highlighting their potential as therapeutic tools for targeting Survivin in cancer treatment.

One of the most significant outcomes of our study was the validation of the deep sequencing-based screening approach. By leveraging this method, we were able to identify seven distinct nanobody clones, all of which showed positive reactivity to Survivin in ELISA assays. This is notable because screening based solely on traditional bio-panning methods often yields a high number of non-specific or low-affinity binders. However, the combination of deep sequencing allowed us to select clones based on both their amplification rate and their abundance in the final pool of binders. This resulted in the enrichment of highly specific nanobodies, all of which demonstrated positive binding in ELISA, indicating that deep sequencing can be a highly efficient and reliable tool for isolating specific, high-affinity nanobodies. In particular, the amplification rate of individual clones proved to be an important metric in selecting optimal binders. While the relative proportion of a clone in the final round of panning could be one factor indicating enrichment, we found that focusing on the amplification rate—how much a clone increases in representation between rounds—was more predictive of superior binding affinity. Clones with higher amplification rates were more likely to show strong binding in subsequent validation assays, suggesting that this metric can be a better criterion for selecting the most effective nanobodies in phage display screenings. Indeed, Clone 7, which demonstrated superior activity across various applications and exhibited a high binding affinity with a Kd of 1.52 µmol/L, also showed the highest amplification factor among the clones, supporting the reliability of using amplification factor as a selection criterion.

Survivin is known to have domain-specific interactions with various partner proteins. Clones 7 and 15 specifically bind to the α-helix region, which is critical for the precise localization of Survivin during mitosis by interacting with proteins in the CPC, such as INCENP (Wheatley and Altieri, 2019). Consistent with this, intracellular expression of these clones suppressed cell proliferation by affecting the cell cycle, raising the possibility that the nanobodies could disrupt Survivin’s interactions with its binding partners, thereby antagonizing its biological functions in the cell. Furthermore, the epitope prediction using AlphaFold Multimer provided valuable structural insights into the binding sites of these nanobodies. The predicted epitopes aligned well with the results of our Western blot analyses, where the nanobodies recognized truncated forms of Survivin containing the alpha-helix region. This consistency between the AlphaFold predictions and experimental data supports the utility of AlphaFold Multimer as a powerful tool for predicting nanobody-antigen interactions. Given its accuracy in predicting binding regions, AlphaFold could be a valuable resource in guiding future nanobody design and in understanding the structural basis of nanobody specificity.

Clone 7 showed a greater effect on reducing cell proliferation compared to Clone 15, likely due to differences in their binding affinity for Survivin. Clone 7 exhibited a stronger binding affinity to Survivin, which may account for its more pronounced inhibitory effect on cell growth. This observation suggests that the binding strength of a nanobody is a key factor in its functional impact, particularly in therapeutic applications where high-affinity binding is crucial for effective inhibition of the target protein’s activity. However, our data also indicated that Clone 15 could achieve a similar level of efficacy when overexpressed using plasmid transfection. This finding highlights the importance of optimizing nanobody expression levels in therapeutic applications, as even nanobodies with lower intrinsic binding affinities can be effective when overexpressed.

In conclusion, this study highlights the utility of deep sequencing as a powerful tool in nanobody screening, enabling the selection of specific high-affinity binders of Survivin with functional relevance. Our findings suggest that Survivin-targeting nanobodies hold considerable potential as therapeutic tools for cancer treatment, particularly in contexts where inhibition of Survivin can disrupt both cell proliferation and survival mechanisms.

## Methods

### Preparation of recombinant antigens

The cDNA of Survivin was amplified by PCR using a cDNA library from HeLa cells reverse-transcribed using oligo dT and SuperScript II (ThermoFisher). For GST-Survivin used in immunization, the cDNA was cloned into the pGEX-6P-1 plasmid, resulting in pGEX-Survivin, which encodes full-length human Survivin fused with GST at the N-terminus. The plasmid was transformed into a chemically competent Rosseta2(DE3)pLysS E.coli strain (Merck Millipore, 71403). The cells were plated on LB agar supplemented with 100 µg/mL Ampicillin and then cultured at 37 °C overnight until colonies were observed. A single colony was inoculated into 2mL of 2YTA (16 g of tryptone, 10 g of yeast extract, and 5 g of NaCl per 1L, containing 100 µg/ml of Ampicillin) and incubated at 37°C overnight with shaking at 200 rpm. 1 ml of the pre-culture was inoculated into 50 ml of 2YTA and cultured at 37°C until A_600_ 0.6. The protein induction was then started by adding 0.1 mM isopropyl β-D-1-thiogalactopyranoside (IPTG). The cultures were grown further with shaking at 200 rpm at 18°C overnight and harvested by centrifugation at 4000×g for 15 min at 4 °C. The cell pellet was stored at -20°C until purification. All purification steps were performed on ice. The E.coli cells were resuspended in 5 ml of Lysis buffer (20 mM HEPES pH8.0, 500 mM NaCl) supplemented with 10 µg/ml DNase I, 1 µg/ml RNase A, 20 µg/ml Lysozyme, 0.2mM PMSF, and 1x CellLytic B Cell Lysis Reagent (Sigma, C8740) and then incubated at 37°C for 5 min. The cell lysate was sonicated using a BioRuptor II (CosmoBio) at high power with 30 seconds ON and 30 seconds OFF cycles for 10 min. The lysate was centrifuged at 15000 rpm for 10 min to remove the insoluble fractions. The cleared lysate was then mixed with 1 ml of glutathione sepharose resin (Cytiva) for affinity purification of GST-tagged survivin. After extensive washing of the resin with PBS supplemented with 0.5 M NaCl, non-tagged Survivin was eluted by on-column cleavage of the GST tag using PreScission protease (Cytiva). pHSM2-Survivin, pHSM-Survivin.Ag1, pHSM-Survivin.Ag2, and pHSM-Survivin.Ag3 were used to express proteins fused with HisTag-SUMO tag at the N-terminus. Expressions using these plasmid DNAs were performed as described above. The expressed proteins were purified by metal affinity chromatography (IMAC) using TALON resin (Takara) as manufacturer’s instructions. The purity of all the purified proteins was analyzed by SDS-PAGE with Coomassie Brilliant Blue (CBB) staining.

### Preparation of recombinant nanobodies

cDNA of each nanobody was cloned into either plasmid pYMVHH7-nH6ND, encoding nanobody with His-bdNEDD8 tag at its N-terminus (Frey and Görlich, 2014b). Two versions of plasmid DNA were prepared: one for expressing the monomeric nanobody, and the other for a tandem dimer nanobody. The plasmids were transformed into the SHuffle *E. coli* strain (NEB, C3030). Induction and purification of HisNEDD8-tagged nanobodies were performed using IMAC, as described above.

### Immunization of survivin to a llama

The immunization of llamas with purified GST-Survivin protein was outsourced to Capralogics (United States, MA). A llama (Llama glama) was injected four times with purified GST-Survivin. 10 days after the final immunization, approximately 5 × 10^8^ peripheral blood mononuclear cells (PBMCs) were isolated from blood samples and suspended in RNAlater (Sigma). The serum and plasma was used for ELISA assays.

### Preparation of a VHH library

Total RNA of PBMC was extracted from the PBMCs using the Sepasol-RNA I SuperG (Nakarai, 09379-84), following the manufacturer’s instructions. cDNA was synthesized by reverse-transcription using PrimeScript II RTase (Takara, 2690A) and an oligo dT primer. VHH cDNA was amplified by nested PCR with two primer sets CALL001:GTCCTGGCTGCTCTTCTACAAGG, CALL002:GGTACGTGCTGTTGAACTGTTCC for the 1st PCR, and VHHYM2_SfiIF2: tgctcctcgcGGCCCAGCCGGCCATGGCTCAGGTGCAGCTGGTGGAGTCTGGGGGAG, VHHYM2-SapIR: GGTCGACGAATTCGGCTCTTCaGCTTGAGGAGACGGTGACCTGGGTCCC for the 2nd PCR. The PCR amplicons were assembled with a phagemid vector pYMPh5.3C_CGGC using T4 DNA ligase (NipponGene, 311-00404). The resulting DNA was transformed to an electro-competent ER2738 E.coli strain (Lucigen, 60522-1) and cultured on 2YTG (2% glucose) plates containing 20 µg/ml of Chloramphenicol. The library size obtained was approximately 3.3 ×10^9^. All colonies were harvested and preserved as 16% glycerol stocks. 2.8 ml of glycerol stock corresponding to 1.6 × 10^11^ cells was inoculated into 1 L of 2YT containing 2% glucose and cultured at 37°C until A_600_ 0.6∼07. M13KO7 helper phage (NEB, N0315S) was infected at a multiplicity of infection (MOI) of 10 for 30 min at RT. After an additional 30 minutes of shaking incubation, the culture medium was replaced with 2YT containing 20 µg/ml chloramphenicol and 50 µg/ml kanamycin, and the culture was continued at 30°C for 19 hours. After centrifugation of the culture medium at 4200 rpm for 30 min, the phage in the supernatant was collected by PEG precipitation. The resulting phage pellet was dissolved in PBS. After measuring the phage titer by qPCR with a primer set StrepT-SeqF01: TGGTCTCACCCGCAGTTCGA and fd_seqR01: GCATTCCACAGACAGCCCTC, the sample was stocked in 8% glycerol at -80°C until use. The display of nanobody protein in the phage was confirmed by western blotting with anti-gpIII antibody.

### Screening of nanobody clones

Screening of nanobodies carried out following previously established protocols (Pardon et al., 2014), with minor adjustments. Briefly, 1 × 10^11^ phages from the library, suspended in 100 µl of PBST containing 0.5% casein, were pre-cleared by incubation with HisSUMO protein conjugated to Dynabeads HisTag (ThermoFisher, 10103D) three times to eliminate anti-SUMO nanobodies. His-SUMO-Survivin coupled with Dynabeads HisTag was used for affinity capture of nanobodies. After blocking the magnetic beads with PBST containing 0.5% casein, the beads were incubated with the pre-cleared phage in a 96 well plate for 2 hours at room temperature. The beads were then washed seven times with PBST, and phages were eluted by treatment with GST-Ulp1 at 30°C for 1 hour. The eluate was inoculated into *E. coli* ER2738 cells (A_600_ 0.5) for 1 hour, followed by infection with M13KO7 helper phage for an additional 1 hour. After changing the culture medium to 2YT containing 20 µg/ml of Chloramphenicol and 50 µg/ml of kanamycin, the culture was grown at 30°C at 1500 rpm overnight in a 96 well deep plate. A total of five rounds of bio-panning were performed, using decreasing amounts of HisSUMO-Survivin protein in each round: 1, 0.2, 0.1, 0.1, and 0.1 µg, respectively. Only for the third round, the antigen was tethered to the bottom of a 96-well Maxisorp plate (ThermoFisher, 442404), conjugated with anti-HisSUMO antibody, instead of using Dynabeads HisTag as in the other rounds, to eliminate non-specific binders to the magnetic beads.

### Alignment and clustering of VHHs

Multiple alignment and clustering of amino acid sequences around the CDR3 region of the top 20 VHH candidates, ranked by the amplification factor, were performed using MAFFT (Katoh, 2002).

### ELISA

For ELISA using the final round of phage displaying nanobody, 10 ng/well of protein (HisSUMO-Survivin, HisSUMO, BSA) was coated onto a 96 well Maxisorp plate through anti-HisSUMO antibody. The wells were blocked with the blocking buffer (2% BSA/PBS) for 1 hour, incubated with 50 µl of phage-containing bacterial culture medium for 1.5 hours at RT, and washed with PBST. Anti-M13 Fd F1 biotin antibody (1:4000) was added for 1 hour, followed by StreptAvidin-HRP (1:4000). After washing, 3,3’,5,5’-Tetramethyl -benzidine (TMB) substrate was applied to detect HRP activity using ABS450 Plate reader (Multiskan FC). For the ELISA using *E. coli* C3030 lysate expressing HisNEDD-nanobody, 40 ng/well of antigen was directly coated onto a 96-well Maxisorp plate, followed by blocking with 2% BSA/PBS. Nanobody was extracted from 1 ml of bacterial culture using 100 µl of lysis buffer (PBS containing 1 mM PMSF, 10% glycerol, 10 µg/ml DNase I, 20 µg/ml lysozyme, and 1x CellLytic B Cell Lysis Reagent). The lysate was diluted 1:300 with the blocking buffer (0.5% BSA/PBS/0.05% Tween20) and incubated with the antigen overnight at 4°C. For detection, StrepTactin-HRP (IBA, 2-1502-001) was used at a 1:2000 dilution. For ELISA using anti-serum, the serum was first pre-cleared six times using Sepharose resin coupled with GST-SUMO to completely deplete anti-GST and anti-SUMO antibodies. The serum, diluted 1:1000 in blocking buffer (0.5% BSA/PBS), was incubated with the antigen, and Goat anti-llama IgG antibody-HRP (BETHYL, A160-100P) was used for detection (1:10000 dilution by blocking buffer).

### Cell culture

mESCs (E14) were cultured in DMEM containing 20% FBS, leukemia inhibitory factor, 3 uM CHIR99021 (a GSK3b inhibitor, Sigma), 1 uM PD0325901 (a MEK inhibitor, Sigma), 1 mM sodium pyruvate, penicillin/streptomycin, non-essential amino acids, and 0.1 mM 2-mercaptoethanol. HEK293T and Hela cells were cultured in IMDM containing 10% FBS and penicillin/streptomycin. The cells were grown at 37°C with 5% CO_2_. For overexpression of FLAG-Survivin, mESCs were transfected with pFL1-hSurvivin using Lipofectamine2000 (ThermoFisher, 11668027).

### Preparation of lentiviruses

Lentiviral vectors pLv-Nb89.7×2-iBSD, pLv-Nb89.9×2-iBSD, pLv-Nb89.15×2-iBSD, and pLv-NbMoonx2-iBSD, encoding nanobodies and a blasticidin resistance gene, were co-transfected into HEK293T cells along with the packaging plasmids pCMV-VSV-G and psPAX2 using FugeneHD (Roche) according to the manufacturer’s instructions. One day after transfection, the medium was replaced, and the cells were cultured for an additional four days. The culture medium was clarified by centrifugation at 15,000 rpm for 15 minutes and stored at -80°C until use.

### Western blotting

Whole cell extracts from the indicated cells were subjected to SDS-PAGE, and the proteins were transferred to a PVDF membrane (Millipore) using the TransBlot Turbo system (Bio-Rad). After blocking with 5% skim milk in PBST, the membranes were incubated with primary antibodies. Tandem dimers of nanobodies were used for all the western blottings. A detailed list of primary and secondary antibodies used is provided in the antibody section.

### Immunostaining

The indicated cells were seeded onto a film-bottom 96-well plate (Greiner 655866) with 1/1000 diluted iMatrix511 (Takara, 892011). After fixation with 4% paraformaldehyde/PBS for 30 minutes at 37°C, cells were treated with 10 mM NH_4_Cl in PBS at room temperature for 10 minutes and then permeabilized with PBS containing 0.1% Triton X-100 for 15 minutes at room temperature. Following a 1-hour block with 2% BSA/PBS, cells were incubated with primary antibodies overnight at 4°C and secondary antibodies for 30 minutes at room temperature. After each antibody treatment, cells were washed three times with PBST and crosslinked again with 4% paraformaldehyde/PBS for 15 minutes at room temperature. Cells were then stained with Propidium Iodide (PI), and fluorescent images were acquired using a Cell Voyager CV1000 spinning disk microscope (Yokogawa) with a 40x dry objective lens. A detailed list of primary and secondary antibodies used is provided in the antibody section. Monomeric nanobodies were used for staining overexpressed Survivin protein, while tandem dimers of nanobodies were used for staining endogenous Survivin protein. For staining with nanobodies, recombinant HisNEDD-nanobodies were first treated with His-NEDP1 to cleave out the HisNEDD tag, resulting in release of nanobodies in a monomeric or tandem form with StrepTagII.

### Prediction of assembled structures using AlphaFold2

Binding site of each nanobody to Survivin protein was predicted using AlphaFold2 with following parameters: --model_preset=multimer, --max_template_date=“2022-12-01”, --use_gpu_relax=false, --num_multimer_predictions_per_model=5, --models_to_relax=“best”.

### Cell proliferation

HeLa cells were transfected with plasmid DNA pNbMoon_iH2BKOP, pNbv89.7_iH2BKOP, or pNbv89.15_iH2BKOP, encoding a monomeric nanobody and puromycin resistance gene, using HilyMax (Dojindo, H357) according to the manufacturer’s instructions. One day after transfection, the cells were cultured with 1 µg/ml of puromycin for 2 days to eliminate non-transfected cells. 1 × 10^5 cells were seeded onto a 24-well plate. After an additional 2 days of culture, the cells were washed twice with PBS, trypsinized to collect adherent cells, stained with Trypan Blue, and counted using a Countess II Automated Cell Counter (Invitrogen).

### Detection of apoptosis

HeLa cells were infected with lentivirus encoding a tandem dimer of nanobody and blasticidin resistant gene. 1 day after the infection, cells were treated with 25 µg/ml of blasticidin for 1 day to eliminate non-infected cells. 5 × 10^4 cells were seeded onto a 12-well plate. After an additional 2 days of culture without any selection reagents, all the cells were collected and stained with Annexin V-FITC Apoptosis Detection Kit (Dojindo) as manufacturer’s instructions with propidium iodide (Dojindo, 341-07881) and DAPI. Fluorescent signals per cell were quantified using Spectral Cell Analyzer SA3800 (SONY).

### Evaluation of CellCycle

After trypsinization, the indicated cells were stained with DAPI in PBS containing 0.1% Triton X-100 for 10 minutes at room temperature. Fluorescent signals for DAPI were then quantified using the Spectral Cell Analyzer SA3800 (SONY). Cell cycle was analyzed based on DAPI intensities.

### Antibodies

Biotin anti-M13 Fd F1 antibody[B62-FE2] (Abcam, ab20337)

Goat Anti-llama IgG antibody-HRP (BETHYL, A160-100P)

StrepTactin-HRP (IBA, 2-1502-001)

StrepTactin Oyster 645 (IBA, 2-1553-050)

anti-Survivin (D-8) (Santa Cruz, sc-17779)

anti-Survivin (FL-142) (Santa Cruz, sc-10811)

anti-α-Tubulin Mouse mAb (DM1A) (Calbiochem, CP06)

anti-FLAG M2 (Sigma, F3165)

anti-FLAG 1E6 (Wako, 014-22383)

anti-Histone H3 (abcam, ab176842)

anti-HisSUMO antibody

## Acknowledgements

Y.M. is supported by JSPS KAKENHI (21H04765, 23K21749, 22H04688), World Premier International Research Center Initiative (WPI, MEXT), Astellas Foundation, Takeda Science Foundation, The Mitsubishi Foundation, The Hokuriku Cancer Foundation, and the Naito Foundation.

## Author contributions

YM. and A.T. designed and performed experiments, and wrote the manuscript. T.F. prepared antigen for immunization.

## Competing interests

All the authors declare no competing interests.

## Notes

### Competing Interest Statement

The authors have declared no competing interest.

## References

Altieri, D.C., 2008. Survivin, cancer networks and pathway-directed drug discovery. Nat. Rev. Cancer 8, 61–70. 10.1038/nrc2293

Ambrosini, G., Adida, C., Altieri, D.C., 1997. A novel anti-apoptosis gene, survivin, expressed in cancer and lymphoma. Nat. Med. 3, 917–921. 10.1038/nm0897-917

Bryant, P., Pozzati, G., Elofsson, A., 2022. Improved prediction of protein-protein interactions using AlphaFold2. Nat. Commun. 13, 1265. 10.1038/s41467-022-28865-w

Cánovas, P.M., 2024. Survivin Mediates Mitotic Onset in HeLa Cells Through Activation of the Cdk1-Cdc25B Axis. Res. Sq. rs.3.rs-3949429. 10.21203/rs.3.rs-3949429/v1

Evans, R., O’Neill, M., Pritzel, A., Antropova, N., Senior, A., Green, T., Žídek, A., Bates, R., Blackwell, S., Yim, J., Ronneberger, O., Bodenstein, S., Zielinski, M., Bridgland, A., Potapenko, A., Cowie, A., Tunyasuvunakool, K., Jain, R., Clancy, E., Kohli, P., Jumper, J., Hassabis, D., 2021. Protein complex prediction with AlphaFold-Multimer. 10.1101/2021.10.04.463034

Fortugno, P., Wall, N.R., Giodini, A., O’Connor, D.S., Plescia, J., Padgett, K.M., Tognin, S., Marchisio, P.C., Altieri, D.C., 2002. Survivin exists in immunochemically distinct subcellular pools and is involved in spindle microtubule function. J. Cell Sci. 115, 575–585. 10.1242/jcs.115.3.575

Frey, S., Görlich, D., 2014a. A new set of highly efficient, tag-cleaving proteases for purifying recombinant proteins. J. Chromatogr. A 1337, 95–105. 10.1016/j.chroma.2014.02.029

Garg, H., Suri, P., Gupta, J.C., Talwar, G.P., Dubey, S., 2016. Survivin: a unique target for tumor therapy. Cancer Cell Int. 16, 49. 10.1186/s12935-016-0326-1

Guo, Y., Mantel, C., Hromas, R.A., Broxmeyer, H.E., 2008. Oct-4 is critical for survival/antiapoptosis of murine embryonic stem cells subjected to stress: effects associated with Stat3/survivin. Stem Cells Dayt. Ohio 26, 30–34. 10.1634/stemcells.2007-0401

Honda, R., Körner, R., Nigg, E.A., 2003. Exploring the Functional Interactions between Aurora B, INCENP, and Survivin in Mitosis. Mol. Biol. Cell 14, 3325–3341. 10.1091/mbc.e02-11-0769

Huang, Y., Fu, J., Zhong, Y., Shuai, W., Zhang, H., Li, Y., He, Q., Tu, Z., 2021. Tandem nanobody: A feasible way to improve the capacity of affinity chromatography. J. Chromatogr. B 1173, 122678. 10.1016/j.jchromb.2021.122678

Jaiswal, P.K., Goel, A., Mittal, R.D., 2015. Survivin: A molecular biomarker in cancer. Indian J. Med. Res. 141, 389–397. 10.4103/0971-5916.159250

Jumper, J., Evans, R., Pritzel, A., Green, T., Figurnov, M., Ronneberger, O., Tunyasuvunakool, K., Bates, R., Žídek, A., Potapenko, A., Bridgland, A., Meyer, C., Kohl, S.A.A., Ballard, A.J., Cowie, A., Romera-Paredes, B., Nikolov, S., Jain, R., Adler, J., Back, T., Petersen, S., Reiman, D., Clancy, E., Zielinski, M., Steinegger, M., Pacholska, M., Berghammer, T., Bodenstein, S., Silver, D., Vinyals, O., Senior, A.W., Kavukcuoglu, K., Kohli, P., Hassabis, D., 2021. Highly accurate protein structure prediction with AlphaFold. Nature 596, 583–589. 10.1038/s41586-021-03819-2

Katoh, K., 2002. MAFFT: a novel method for rapid multiple sequence alignment based on fast Fourier transform. Nucleic Acids Res. 30, 3059–3066. 10.1093/nar/gkf436

Li, F., Aljahdali, I., Ling, X., 2019. Cancer therapeutics using survivin BIRC5 as a target: what can we do after over two decades of study? J. Exp. Clin. Cancer Res. 38, 368. 10.1186/s13046-019-1362-1

Li, Y., Lu, W., Yang, J., Edwards, M., Jiang, S., 2021. Survivin as a biological biomarker for diagnosis and therapy. Expert Opin. Biol. Ther. 21, 1429–1441. 10.1080/14712598.2021.1918672

Lobstein, J., Emrich, C.A., Jeans, C., Faulkner, M., Riggs, P., Berkmen, M., 2012. SHuffle, a novel Escherichia coli protein expression strain capable of correctly folding disulfide bonded proteins in its cytoplasm. Microb. Cell Factories 11, 753. 10.1186/1475-2859-11-56

Lv, Y.-G., Yu, F., Yao, Q., Chen, J.-H., Wang, L., 2010. The role of survivin in diagnosis, prognosis and treatment of breast cancer. J. Thorac. Dis. 2, 100–110.

Olie, R.A., Simões-Wüst, A.P., Baumann, B., Leech, S.H., Fabbro, D., Stahel, R.A., Zangemeister-Wittke, U., 2000. A novel antisense oligonucleotide targeting survivin expression induces apoptosis and sensitizes lung cancer cells to chemotherapy. Cancer Res. 60, 2805–2809.

Pardon, E., Laeremans, T., Triest, S., Rasmussen, S.G.F., Wohlkönig, A., Ruf, A., Muyldermans, S., Hol, W.G.J., Kobilka, B.K., Steyaert, J., 2014. A general protocol for the generation of Nanobodies for structural biology. Nat. Protoc. 9, 674–693. 10.1038/nprot.2014.039

Shin, S., Sung, B.-J., Cho, Y.-S., Kim, H.-J., Ha, N.-C., Hwang, J.-I., Chung, C.-W., Jung, Y.-K., Oh, B.-H., 2001. An Anti-apoptotic Protein Human Survivin Is a Direct Inhibitor of Caspase-3 and -7. Biochemistry 40, 1117–1123. 10.1021/bi001603q

Stauber, R.H., Mann, W., Knauer, S.K., 2007. Nuclear and cytoplasmic survivin: molecular mechanism, prognostic, and therapeutic potential. Cancer Res. 67, 5999–6002. 10.1158/0008-5472.CAN-07-0494

Sui, L., Dong, Y., Ohno, M., Watanabe, Y., Sugimoto, K., Tokuda, M., 2002. Survivin expression and its correlation with cell proliferation and prognosis in epithelial ovarian tumors. Int. J. Oncol. 10.3892/ijo.21.2.315

Temme, A., Rieger, M., Reber, F., Lindemann, D., Weigle, B., Diestelkoetter-Bachert, P., Ehninger, G., Tatsuka, M., Terada, Y., Rieber, E.P., 2003. Localization, Dynamics, and Function of Survivin Revealed by Expression of Functional SurvivinDsRed Fusion Proteins in the Living Cell. Mol. Biol. Cell 14, 78–92. 10.1091/mbc.e02-04-0182

Uren, A.G., Wong, L., Pakusch, M., Fowler, K.J., Burrows, F.J., Vaux, D.L., Choo, K.H.A., 2000. Survivin and the inner centromere protein INCENP show similar cell-cycle localization and gene knockout phenotype. Curr. Biol. 10, 1319–1328. 10.1016/S0960-9822(00)00769-7

Vader, G., Kauw, J.J.W., Medema, R.H., Lens, S.M.A., 2006. Survivin mediates targeting of the chromosomal passenger complex to the centromere and midbody. EMBO Rep. 7, 85–92. 10.1038/sj.embor.7400562

Wheatley, S., Mcneish, I., 2005. Survivin: A Protein with Dual Roles in Mitosis and Apoptosis. Int. Rev. Cytol. 247, 35–88. 10.1016/S0074-7696(05)47002-3

Wheatley, S.P., Altieri, D.C., 2019. Survivin at a glance. J. Cell Sci. 132, jcs223826. 10.1242/jcs.223826

Wheatley, S.P., Carvalho, A., Vagnarelli, P., Earnshaw, W.C., 2001. INCENP is required for proper targeting of Survivin to the centromeres and the anaphase spindle during mitosis. Curr. Biol. 11, 886–890. 10.1016/S0960-9822(01)00238-X

